# Global Analyses of Genomic and Epigenomic Influences on Gene Expression Reveals *Serpina3n* as a Major Regulator of Cardiac Gene Expression in Response to Catecholamine Challenge During Heart Failure

**DOI:** 10.1101/2025.09.07.674751

**Authors:** Caitlin Lahue, Sriram Ravindran, Aryan Dalal, Rozeta Avetisyan, Christoph D Rau

## Abstract

Heart failure arises from maladaptive remodeling driven by genetic and epigenetic networks. Using a systems genetics framework, we mapped how DNA variants and CpG methylation shape cardiac transcriptomes during beta adrenergic stress in the Hybrid Mouse Diversity Panel, a cohort of over 100 fully inbred mouse strains. Expression QTLs (eQTLs), methylation QTLs (mQTLs) and methylation-driven eQTLs (emQTLs) were generated from over 13k expressed genes and 200k hypervariable CpGs in left ventricles. We discovered hundreds of regulatory “hotspots” that control large portions of the genome, including several that regulate over 10% of the transcriptome and/or methylome. Approximately 16% of these hotspots overlapped with prior GWAS or EWAS signals. We focus on a hotspot on chromosome 12 and identify the serpine peptidase inhibitor *Serpina3n*, as the most likely driver gene in this hotspot. Experimental knockdown of *Serpina3n* in neonatal rat ventricular cardiomyocytes blunted hypertrophy induced by a variety of hypertrophic signals, while altering predicted target expression and modulating the activity of *Nppa* and *Nppb*. Together, these findings position *Serpina3n* as a major regulator of stress-responsive cardiac gene programs, highlighting how integration of genetic and epigenetic signals can pinpoint key drivers of heart failure.

**Key Highlights:**

- **Cross-omics hotspot analysis**: We identified 286 eQTL, mQTL, and emQTL hotspots in the Hybrid Mouse Diversity Panel, including several hotspots that regulate over 10% of the cardiac transcriptome.
- **Integration with prior GWAS/EWAS**: ∼16% of hotspots overlapped with previously reported heart failure hotspots, strengthening their biological relevance.
- **Discovery of Serpina3n as heart failure regulator**: A hotspot on chromosome 12 linked to *Serpina3n* controlled 4–6% of all gene expression and CpG methylation changes in response to isoproterenol.
- **Functional validation**: siRNA knockdown of *Serpina3n* in cardiomyocytes significantly blunted hypertrophy induced by isoproterenol, angiotensin II, and phenylephrine.
- **Pathway insights**: Genes regulated by the *Serpina3n* hotspot were enriched for mitochondrial function, dilated cardiomyopathy, and hypertrophic signaling pathways, highlighting a mechanistic link to heart failure progression.

## Introduction

Heart failure is a major driver of mortality increasing health care costs. Heart failure’s causes are diverse, multi-faceted and characterized by the interaction of complex transcriptional networks with external stimuli that act together to steer the heart from a healthy to diseased state^1^. Significant research effort has been directed towards understanding the underlying genetic and epigenetic changes that contribute to heart failure, including genome and epigenome-wide association studies (GWAS/EWAS)^2–7^. Despite identifying a number of interesting loci and genes for future study, the link between genetic/epigenetic variation and the underlying transcriptional networks which mediate the development of heart failure remains incompletely understood. The majority of GWAS loci for complex traits are thought to act through regulation of gene expression, and it is broadly accepted that changes in DNA methylation are inversely correlated to the expression of nearby genes^8^. Consequently, better understanding how individual polymorphisms and/or shifts in CpG methylation status regulate the expression of genes in the transcriptome in the context of the stressed heart will shed new light on genes and pathways which drive the progression of heart failure and other cardiovascular disorders.

We have previously published a set of studies examining genetic^9,10^ and epigenetic^7^ loci that drive heart-failure-associated phenotypes in the Hybrid Mouse Diversity Panel (HMDP), a murine genetic reference population of over 100 commercially available strains of mice that has also been used to map genetic and epigenetic loci for traits related to high fat diet^11^, diabetes^12^, immune cell abundances in blood^13^, and more^12,14–17^. In our heart failure study, we used chronic administration of the beta adrenergic agonist Isoproterenol (ISO) to drive cardiac hypertrophy and heart failure in our mice. We observed dozens of significant GWAS and EWAS loci using this approach, and identified several novel genes contributing to hypertrophy, cardiac fibrosis, and echocardiographic parameters^4,7,10^.

We now report a detailed analysis of the effects of genetic polymorphisms and CpG methylation variation on gene expression during the progression of heart failure. We performed GWAS on gene expression from over 13,000 expressed genes in the left ventricles of the heart to generate expression Quantitative Trait Loci (eQTLs) and EWAS on gene expression to generate what we term expression methylation-driven Quantitative Trait Loci (emQTLs). We also performed GWAS on CpG methylation sites to generate methylation QTLs (mQTLs). In particular, we focus on the identification of major transcriptional or methylation regulators, or eQTL/mQTL/emQTL hotspots, in which a single SNP or CpG regulates a significantly large number of genes’ expression profiles or a single SNP regulates a large number of changes in CpG methylation statuses across the genome.

eQTL hotspot analysis reveals 49 significant hotspots, including 10 hotspots which regulate over 10% of the genes of the heart. mQTL hotspot analysis returns 98 hotspots, although these are generally smaller with only 2 hotspots regulating over 10% of CpG sites. Finally, emQTL hotspot analysis shows a striking 139 significant hotspots, yet only one regulates over 10% of the genes of the heart. Within these hotspots, we have identified candidate genes with known roles in cardiac growth and the response to adrenergic stimulation, such as *Phlllp1*^18,19^ and *Kcnj2*^20,21^ among other novel candidates. In particular, we focus on the *Serpina3n* gene, which was recently reported to play a major role in the response to myocardial infarction^22^. We shed additional light on its important role in the progression of heart failure, showing that it resides in a eQTL hotspot that regulates the change of expression of 5.6% of all genes (P=3.1*10^-10^) and a emQTL hotspot that regulates the change of expression of 4% of all genes (P=8.5*10^-37^). We further demonstrate that knockout of *Serpina3n* not only reduces cardiac hypertrophy, but directly regulates predicted gene targets as well as hypertrophic markers.

## Methods

### Ethics Statement

All animal experiments were conducted following guidelines established and approved by the University of North Carolina at Chapel Hill Institutional Animal Care and Use Committee. All surgery was performed under isoflurane anesthesia and every effort was made to minimize suffering. Humane endpoints included loss of weight, lethargy, hunched posture and other standard endpoints. Animals were kept in standard cages within the vivarium and provided standard chow and enrichment. We have adhered to the ARRIVE guidelines for this study.

### Hybrid Mouse Diversity Panel Isoproterenol Study

We have previously reported genetic^4,10,23^ and epigenetic^7^ studies of heart failure in the Hybrid Mouse Diversity Panel. 8-10 week old (average 9.1 weeks) female mice from 101 diverse inbred mouse strains were separated into isoproterenol-treated (4 mice per strain) and control (2 mice per strain) cohorts. Isoproterenol (ISO) was administered through an intraperitoneally-implanted osmotic micropump (Alzet, #2004) at a dose of 30 mg ISO/kg body weight/ day. Mice underwent echocardiography before surgery and weekly thereafter. After 21 days, all mice were euthanized, organs collected, weighed, and sectioned for flash freezing in liquid nitrogen and for histology. All mice were obtained from Jackson Labs (Bar Harbor, ME) or directly from the UCLA HMDP colony as previously described^4^. Our study was limited to female mice for cost reasons but an earlier pilot study showed stronger effects on cardiac phenotypes by ISO in female mice and prior work in the HMDP suggests that most results are translatable between sexes^12,15^.

RNA was previously isolated and microarrays generated^4^. To briefly summarize: RNA was isolated from a portion of the left ventricle of two ISO-treated and two control mice per strain for 99 strains via RNeasy kit (Qiagen), tested for RIN > 7 by Agilent Bioanalyzer, and pooled together for one control and one treated sample per strain. RNA abundance was determined via an Illumina Mouse Reference 8 version 2.0 chip. Analysis was performed using the Neqc algorithm in the limma R package^24^ after batch effect correction via COMBAT^25^.

DNA was previously isolated and RRBS performed^7^. To briefly summarize: DNA was isolated from a portion of the left ventricle of one ISO-treated and one control mouse per strain for 90 HMDP strains via dnEasy kit (Qiagen) according to manufacturer instructions and quantified using the Qubit dsDNA HS kit. RRBS was performed as described^26^ in which DNA was digested with MspI, then library prep performed using the NEBNext Ultra DNA library prep kit (NEB E7370L) followed by bisulfite conversion and sequenced using an Illumina HiSeq2500 at 1×101 bp read length. RRBS reads were aligned to the mm10 genome and quantified as described^7^ using the BSseeker2 algorithm^27^ with default parameters. CpGs with Q>=30 and read depth of >3 were preserved and CpGs in strains which had a detected SNP at the CpG site were removed before batch correction with COMBATseq^28^. For the purposes of this manuscript, we focus exclusively on cytosines in CG dinucleotides.

All strains used for genomic, transcriptomic or epigenomic analyses can be found in Table S1.

### eQTL, mQTL, and emQTL generation

Genes for QTL analysis were selected based on transcripts which were expressed (average expression value > 5) and varying (coefficient of variation > 10%) across the HMDP panel in the LV transcriptomes, resulting in a final set of 13,155 genes.

SNPs were selected for eQTL/mQTL analysis based on a minor allele frequency of 5% in the used HMDP strains based on the genotypes generated and used in our prior studies^29^.

CpGs were selected for mQTL/emQTL analysis based on hypervariability as described in earlier methylation studies in the HMDP^16,17,30^. Hypervariable CpGs are defined as CpGs in which the percent methylation shifts by over 25% in at least 5% of the affected strains (5% chosen to mirror a 5% SNP minor allele frequency cutoff).

eQTLs, mQTLs, and emQTLs were generated from these data as follows:

*eQTLs and mQTLs*: Genetic control of population structure for gene expression or DNA methylation was conducted as previously described^10^using the Efficient Mixed Model Algorithm (EMMA), which models the effects of a SNP on a phenotype y through the linear mixed model 𝒚 = 𝒎 + 𝒙 + 𝒖 + 𝒆 where m is the mean phenotypic value, b is the allele effect of the SNP, x is the vector of genotypes for every strain, u is the random effects of genetic relatedness where *var*(*u*) = σ_*u*_^2^*K* with K denoting the kinship matrix relating strains to one another and *var*(*e*) = σ_*e*_^2^*I* with both 𝜎_*u*_^2^ and 𝜎_*e*_^2^ estimated using restricted maximum likelihood.

*emQTLs*: epigenetic control of gene expression was conducted as previously described^30^ through the methylation-specific binomial mixed model approach MACAU^31^. Briefly, MACAU models each CpG as 𝑦_𝑖_∼𝐵(𝑣_𝑖_, 𝜋_𝑖_), with 𝑣_𝑖_ representing the total read count for the ith individual, 𝑦_𝑖_ as the methylated read count (with 𝑦_𝑖_ ≤ 𝑣_𝑖_) and 𝜋_𝑖_ denoting an unknown parameter that is the true proportion of methylated reads at that CpG. MACAU then models 𝜋_𝑖_ as follows using a logit link:

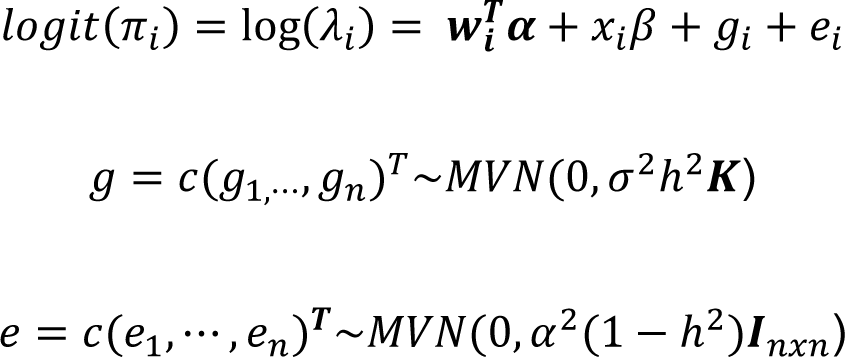

Where 𝒘_𝒊_ is a c-vector of covariates including an intercept and **α** is a c-vector of corresponding coefficients, 𝑥_𝑖_ is the predictor of interest and β is its coefficient. g is a n-vector of genetic random effects that model correlation due to population structure and e is a n-vector of environmental residual errors that model independent variation. **K** is a known n x n relatedness matrix based, in our case, on genotype data, and standardized to ensure that 𝑙𝑣(𝐾)/𝐵 = 1 (this ensures that ℎ^2^ lies between 0 and 1 and can be interpreted as heritability). **I** is a n x n identity matrix, 𝜎^2^ℎ^2^ is the genetic variance component, 𝜎^2^(1 − ℎ^2^) is the environmental variance component, ℎ^2^ is the heritiability of the logit transformed methylation proportion (aka *logit*(𝜋)) and MVN denotes the multivariate normal distribution. MACAU then tests for an association between a CpG and a gene’s expression by testing the null hypothesis (𝛽 = 0). It samples to compute an approximate maximum likelihood estimate 𝛽^, its standard error 𝑠𝑒(𝛽^), and a corresponding p-value of significance as described^31^.

### Hotspot Analysis

The genome/epigenome was binned into windows of 500kb. For each gene/CpG, the lowest p-value within each window was used to represent the relationship of the gene/CpG to SNPs or CpGs within that window as a whole. Hotspot strengths were determined by summing up the number of genes/CpGs within each windows whose lowest p-value in that window was less than 4.2*10^-5^ (a suggestive cutoff approximately one order of magnitude less stringent than the accepted 4.2*10^-6^ genome-wide significance threshold of the HMDP as a whole^32^). Hotspot significance was determined by z-transformation with the significance threshold set at 9.3*10^-6^, which is the Bonferroni-corrected threshold with α=0.05 and 5400 windows (2.7 Gbase / 500 kbase).

### Candidate Gene Selection

All genes within 1 MB in either direction of a hotspot window were considered to account for the average size of linkage disequilibrium in the HMDP^15^. We then leveraged data from other HMDP studies^4,15,33^ to prioritize candidates. First, we looked for exonic variants in the sequenced founders of the HMDP^34^ that were predicted to cause changes in gene expression or function. Next, we examined whether the gene’s expression was correlated with cardiac phenotypes, or whether the gene was associated with cardiac-related phenotypes in prior HMDP studies by GWAS, EWAS, etc. Finally, we explored the literature for published evidence on the gene.

### In Vitro Validation Studies

Neonatal Rat Ventricular Cardiomyocytes (NRVMs) were isolated from 1-4 day old Wistar rats as previously described^30^ using the Cellutron Neomyocyte isolation kit. Briefly, hearts were excised, washed in PBS and trimmed, then placed in ice cold PBS. After 10 hearts were isolated, PBS was removed and replaced with 4 mL Cellutron Digestion Buffer, then incubated for 12 minutes at 37 C on a stir plate (150 rpms) in a 25 mL beaker with a 1 inch stir bar.

Supernatant containing isolated cells were transferred to a new 15 mL tube and spun at 2200 rpm for 2 minutes. Supernatant was discarded and cells resuspended in digestion stop buffer supplemented with cell media and stored. Simultaneously, 4 mL of digestion buffer had been added to the hearts and the entire process repeated approximately 8 times until the heart turned pale pink and notably fewer cells were recovered after centrifugation. All 15 mL tubes were spun down, supernatants discarded, and final cells resuspended in 2 mL ADS buffer (12mM NaCl, 2mM HEPES, 1 mM NaH2PO4, 0.5mM Glucose, 0.5mM KCl, 0.1 mM MgSO4).

NRVMs were purified using a Percoll gradient. 6 mL of 1.059g/mL Percoll was layered atop 3 mL of 1.082g/mL Percoll diluted in ADS buffer in a 15 mL conical tube. 2 mL cell suspension was layered on top of the Percoll mixtures and the entire tube centrifuged at 3000 rpm for 30 minutes at the slowest possible ramp up speed and with no brake. After centrifugation, the lower band of cells containing NRVMs was extracted and diluted in 10 mL of ADS buffer followed by centrifugation at 2200 rpm for 10 minutes. Cell pellet was resuspended in 2 mL DMEM + 10% PBS and 1% pen/strep, followed by counting on a Biorad TC20 cell counter (ThermoFisher) and plating onto gelatin-coated 12-well plates at a density of 200-250k cells/well.

We followed our previously established protocol for testing gene siRNAs in NRVMs (see Table S2 for siRNAs). 24 hours after plating, DMEM media containing FBS and pen/strep was aspirated and wells washed 2x in PBS. DMEM media containing 1% ITS supplement (SigmaAldrich) was added to each well. That same day, siRNAs were transfected into cells using lipofectamine RNAiMax (Invitrogen) per manufacturer instructions. For each siRNA experiment, 6 wells across 2 12-well plates each got either control (no siRNA), scramble siRNA, or a siRNA obtained from IDT (See Table S2). Transfections were allowed to proceed for 24 hours, then the media was refreshed and isoproterenol added to half of the wells at a final concentration of 10 mM. After 48 hours, photographs of each well were taken at 20x magnification and RNA isolated for qPCR validation of gene knockdown and expression of any correlated gene targets(see Table S2 for all primers). Cell cross-sectional area and confluence were assessed for each well by trained users blinded to the sample identity. Differences between conditions are tested using two-way ANOVA followed by a Tukey’s Range Test for each comparison.

### Gene Ontology Enrichment

Gene ontology enrichment was performed using the Gene Analytics Suite^35^ which uses a binomial test to test the null hypothesis that a defined set of genes is not over-represented within a given pathway and then corrected using the Benjamini-Hochberg correction (FDR). GeneAnalytics has several modules (e.g. a Gene Pathways module, a GO Terms module, etc.) We specify which module we use in the text as needed. All p values reported are corrected p values.

## Results

### Hybrid Mouse Diversity Panel

In prior work^4,10^, we described the study of a cohort of 101 strains of the Hybrid Mouse Diversity Panel (HMDP) for Heart Failure associated traits. 8-10 week old mice were divided into treatment and control cohorts, and treated mice were given the beta adrenergic agonist Isoproterenol (ISO) via an Alzet osmotic pump (at 30 mg/kg body weight/day) for three weeks to cause adrenergic overdrive, cardiac hypertrophy and, in some cases, cardiac failure as measured by echocardiogram. We observed significant variability in the phenotypes of the heart, from changes in cardiac fibrosis (Figure 1A), to fractional shortening (Figure 1B), to other phenotypes in response to ISO. As part of this study, we generated transcriptomes for 99 strains of mice both with and without ISO stimulation, which we used to help narrow down GWAS loci and identify candidate genes. More recently, we have reported^7^ the addition of DNA methylation data generated by Reduced Representational Bisulfite Sequencing (RRBS) from the hearts of 90 strains of the same HMPD HF study. In this study, we calculate expression Quantitative Trait Loci (QTLs) either at the SNP (eQTL) or CpG level (emQTL) to identify whether there are SNPs or CpGs which significantly affect the expression of a large number of genes across the genome (Figure 1C), also known as a ‘hotspot.’ We additionally calculated whether there are methylation QTL (mQTL) hotspots in the genome where a SNP regulates a large number of CpG methylation statuses.

**Figure 1:**
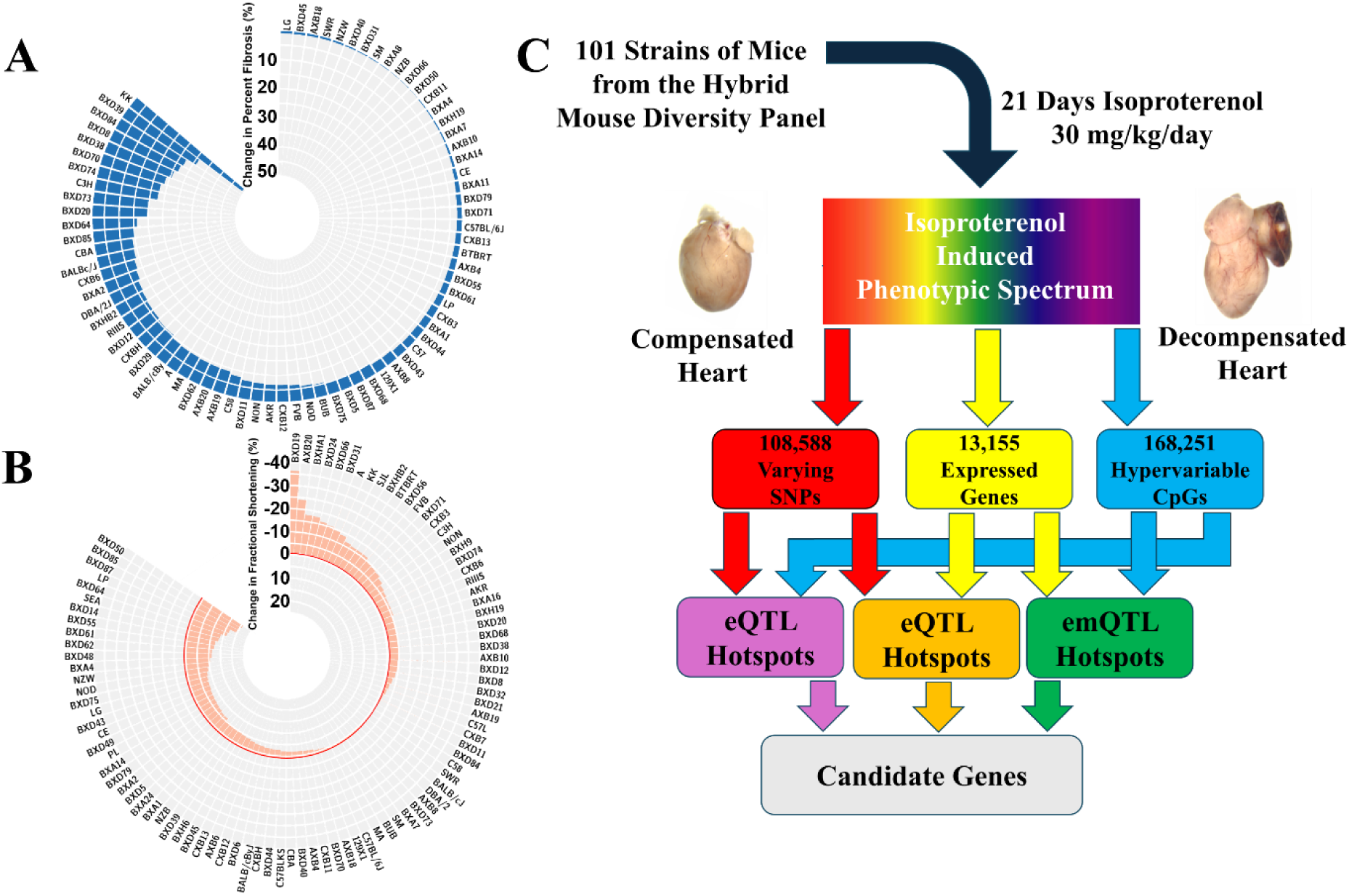
The Hybrid Mouse Diversity Panel Heart Failure Study. A + B) representative circle plots showing the average change in fibrosis (A) or fractional shortening (B) across the HMDP cohort in response to chronic isoproterenol challenge. C) Schematic for the current study, in which SNP, expression, and methylation information was combined together to identify eQTL, mQTL, and emQTL hotspots

### Hotspot Identification

Our data contained 13,155 expressed and variable genes in either the control or ISO conditions (8,126 genes varied in both conditions). We further observe 108,588 hypervariable CpGs across our cohort. We calculated eQTLs and mQTLs in control, treated, and delta (ISO-control) conditions using EMMA^36^ and emQTLs with the MACAU algorithm^31^. The genome was binned into 500kb windows and, for each type of QTL, genes/CpGs with suggestive (P=4.2*10^-5^, one order of magnitude less than the genome-wide significance threshold in the HMPD of 4.2*10^-6^) significances were summed up for each region. Regions that regulated a statistically significant number of genes/CpGs were identified by converting hotspot values to z-scores relative to the null distribution and applying a Bonferroni-adjusted significance threshold of P<9.3*10^-6^.

As shown in Table 1, depending on the condition (Control vs ISO vs Delta) or class (eQTL vs mQTL vs emQTL), we observe between six large hotspots for control eQTL (average size of hotspot over 4 Mb) to forty small hotspots for Isoproterenol mQTL (average size of hotspot 620 Kb) and regulating anywhere from 0.8% of the expressed genes in Isoproterenol emQTLs to 25.1% of changing genes in Delta eQTLs. We observe that our Delta hotspots always seem to regulate more genes than our control hotspots and that our ISO hotspots regulate the least, with the exception of ISO mQTLs, which are the most prominent in their class. (Figure 2). Overall, we observe 286 unique hotspots, with 52 of these hotspots present in multiple classes (Table S3).

**Table 1.** Significant eQTL, mQTL and emQTL Hotspots

| **Condition** | **Class** | **Number of Loci** | **Average Locus Size (Kb)** | **% Genes Regulated** |
| --- | --- | --- | --- | --- |
| Control | eQTL | 6 | 4080 | 2.3 - 13.1 |
| Isoproterenol | eQTL | 16 | 750 | 1.4 - 9.8 |
| Delta | eQTL | 27 | 1090 | 4.0 - 25.1 |
| Control | mQTL | 23 | 870 | 0.9 - 3.1 |
| Isoproterenol | mQTL | 40 | 620 | 4.3 - 11.5 |
| Delta | mQTL | 35 | 700 | 1.5 - 5.2 |
| Control | emQTL | 41 | 1020 | 1.0 - 6.3 |
| Isoproterenol | emQTL | 39 | 1040 | 0.8 - 3.2 |
| Delta | emQTL | 59 | 790 | 2.9 - 12.8 |

**Figure 2:**
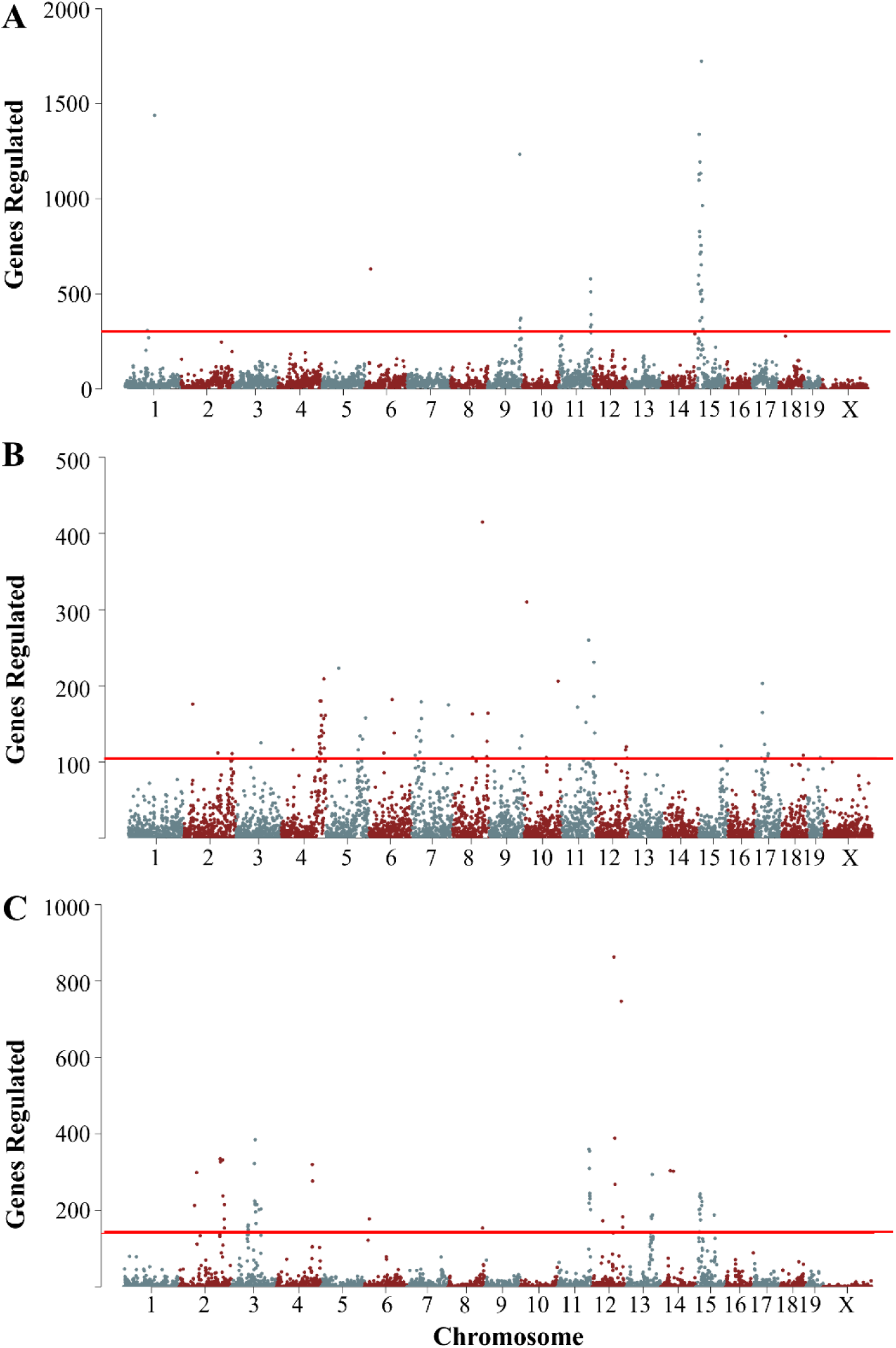
Representative Hotspots for Expression and Methylation. A) Control eQTL hotspots B) ISO emQTL hotspots C) Delta eQTL hotspots. In all cases, points represent the number of genes/CpGs regulated at a suggestive (P=4.2*10^-5^) level within a 500kb window. The red line indicates a genome-wide significance threshold.

### Overlap with prior GWAS/EWAS loci from the HMDP Study

32 (15.7%) of our hotspots overlap with GWAS or EWAS hits from our prior analyses, significantly more than expected by chance (P=1.0*10^-9^). These hotspots include a hotspot on chromosome 7 (Delta_emQTL_24) in which a change in DNA methylation regulates the change of 440 genes across the genome. This hotspot is also a locus for cardiac fibrosis after isoproterenol challenge from our past work and includes the gene *Abcc6*, which we validated *in vivo* as a powerful driver of fibrosis only under the condition of beta adrenergic overdrive^4^.

Similarly, we see an overlap between a CpG hotspot on chromosome 17 (Delta_emQTL_54) that regulates 5.6% of the change of gene expression after ISO challenge in the heart and an EWAS locus for posterior wall thickening that includes *Anks1,* whose broader family of Ankyrins have been implicated in cardiomyopathies, even though the role of *Anks1,* specifically, remains unclear^37^. We also highlight *Slit2*, the candidate gene for the fifth most significant ISO emQTL hotspot (Iso_emQTL_9) which we showed in our EWAS study promoted cardiac growth^38^ See Table S4 for all observed overlaps.

### Novel High-Significance Heart Failure Associated Hotspots

We now turn our focus to hotspots which significantly affect a large percentage of expressed and varying genes or CpGs in the heart, focusing in particular on the 5 most significant hotspots for Control, Isoproterenol, and Delta conditions for eQTLs, mQTLs, and emQTLs (45 total peaks, 39 unique, Table 2). The most striking of these hotspots is a hotspot on chromosome 1 for delta eQTL which regulates just over 25% of the *change* in gene expression in response to ISO in the heart (P=3.8*10^-169^). Of the ten genes present within this hotspot, two have features (in the form of either a significant *cis* eQTL, correlation to important HF-associated phenotypes, or the presence of a protein-altering mutation in one of the five parental lines of the recombinant inbred crosses of the HMDP) that suggest they may be the underlying driving gene of the hotspot. The first is *Pign*, which has a mutation in A, Balb, and C3H. *Pign* is a biosynthesis protein involved in the creation of the GPI anchor glycoprotein, and its loss is causal for multiple congenital anomalies–hypotonia–seizures syndrome 1 (MCAHS1), which has reported sporadic cardiac complications^39^. A stronger candidate based on the literature is the phosphatase *Phlpp1*, which has a mutation in DBA2, acts to dephosphorylate both AKT and PKC, and has been linked to cardiomyocyte death and cardiac dysfunction^19^ and whose inhibition by Isoquercitrin reduces cardiomyocyte apoptosis^18^.

**Table 2.**
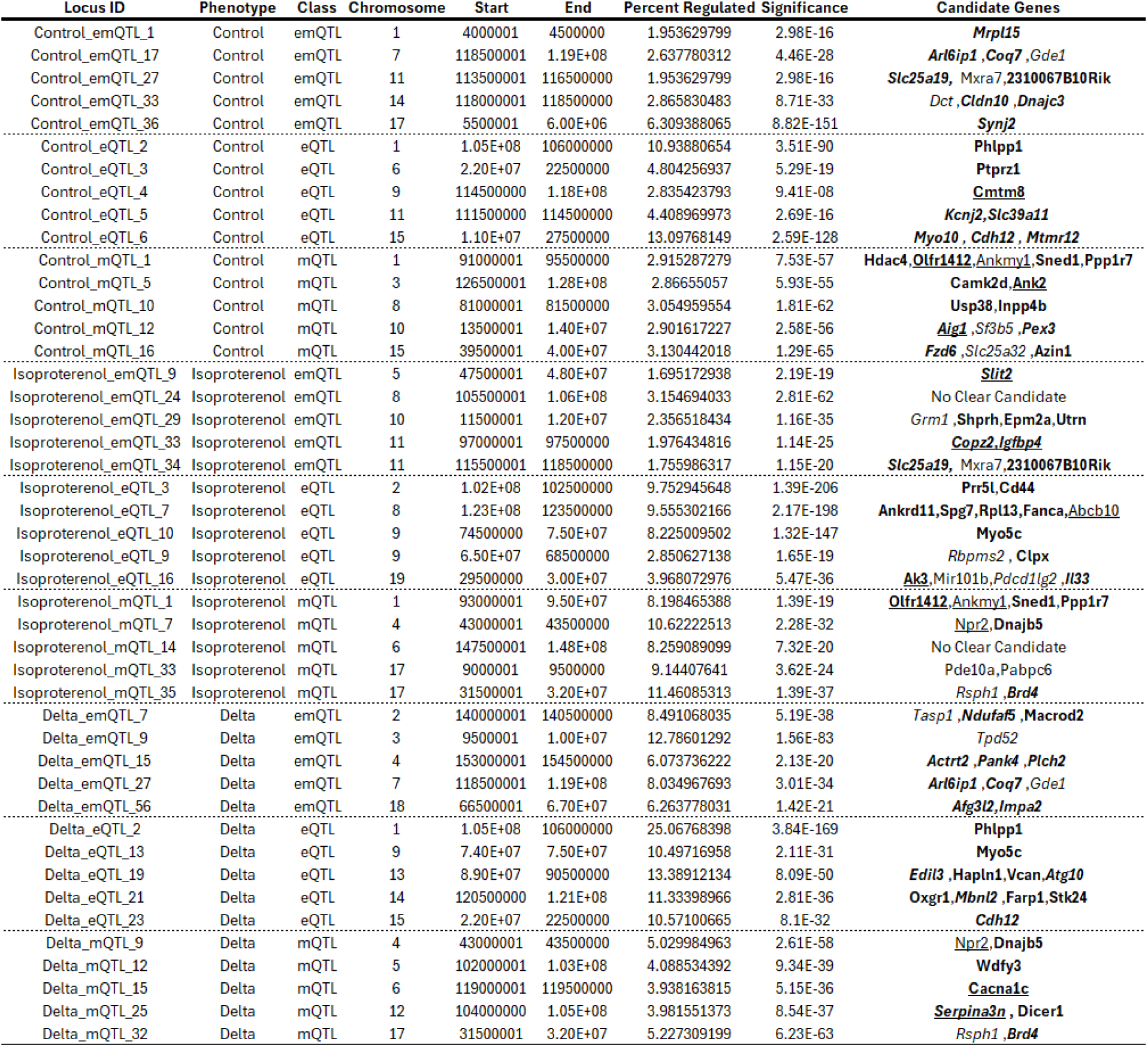
Top 5 hotspots per class and phenotype. . **Bold** entries indicate genes with missense or nonsense mutations in a HMDP founder (A, B6, C3H, DBA). *Italic* entries have a significant cis-eQTL at this hotspot. <u>Underlined</u> entries indicate a significant correlation with a HF-associated trait (e.g. heart weight). See Table S3 for a full list of all hotspots

Another significant hotspot lies on chromosome 8, and controls the expression of 9.6% of all ISO-expressed genes (P=2.2*10^-198^). Curiously, this hotspot also overlaps with a more modest but still significant epigenetic hotspot that controls 1.6% of the expression of *control*-expressed genes (P=1*10^-6^, see Tables S3), suggesting that there are possibly two signals at this location, one driven by genetics and one by epigenetics in distinct contexts. This locus contains several interesting genes, with four genes (*Ankrd11, Spg7, Rpl13* and *Fanca*) each possessing nonsynonymous mutations in one of the HMDP Recombinant Inbred founders and another, *Abcb10,* showing correlation with ISO-treated heart weights (R=0.33,P= 1.3*10^-3^) and a suggestive but not significant cis-eQTL(P=0.018). Both *Ankrd11*, causal for KBG syndrome, and *Rpl13* are linked to congenital heart defects^40,41^, while modulation of *Abcb10* in adult cardiomyocytes led to increased levels of oxidative stress^42^ and broader cardiac dysfunction^43^.

We next turned our attention to hotspots which showed evidence of concurrent regulation across multiple ‘omics layers. On chromosome 11, there is an eQTL/emQTL hotspot that controls 4.4% of all untreated gene expression (P=2.7*10^-16^) and spans 3 Mb from 111.5-114.5Mb. There are two notable candidate genes in this hotspot. *Slc39a11* is a solute carrier with a predicted deleterious nonsynonymous mutation in A and C3H backgrounds and a significant *cis-*eQTL, although its mechanical link to cardiac function is unclear, while the potassium transporter *Kcnj2* shows a strong *cis-eQTL* and is the causal gene for Andersen-Tawil syndrome which presents with, among other phenotypes, dilated cardiomyopathy and additional cardiac dysfunctions^20,21^.

A hotspot which shows even stronger evidence of concurrent trans-omic regulation lies on chromosome 9 and is significant in all three levels of our analysis. At the genetic level, it regulates 373 genes (2.8%, P=9.4*10^-8^) and 2,116 CpGs (1.26%, P=7.2*10^-11^), while at the epigenetic level, it regulates 189 genes (1.43%, P=1.3*10^-9^). This hotspot contains three genes of further interest. The first is *Tgfbr2*, which has a wide variety of roles^44^, including several related to cardiac development and function^45,46^. Despite these observed roles, we see no evidence in the form of significant cis-eQTLs, notable sequence mutations, or correlation to phenotypes to recommend it as the causal gene at this hotspot, suggesting that other genes deserve more scrutiny. The second gene of interest is *Dync1li1*, a dynein protein associated with angiogenesis^47^ and with a strikingly significant correlation with baseline cardiac fibrosis (R=-0.49, P=8*10^-11^) and a strong cis eQTL. The final gene of interest at this hotspot is *Cmtm8*, a putative tumor suppressor and growth modulator, which has a deleterious mutation in the A, Balb, and C3H backgrounds. Curiously, although this gene is correlated with control phenotypes, notably atrial weight (R=0.277,P=0.01) and intraventricular septal width (R=0.290,P=5.2*10^-4^), we observe much stronger correlations of baseline *Cmtm8* gene expression with ISO-treated phenotypes including total heart weight (R=-0.252,P=0.0155), lung and liver weight (R=-0.357,P=4.8*10^-4^, R=-0.380,P=1.9*10^-4^), and ejection fraction (R=0.40,P=8.5*10^-5^). This mirrors what we have previously observed in our EWAS study^30^, where control gene methylation/expression was at times more predictive of ISO phenotypes than the gene’s expression/methylation after ISO.

### Serpina3n is a major regulator of cardiac transcription, methylation, and hypertrophy

For a deeper analysis of an individual hotspot, we selected a hotspot on chromosome 12 from 104-104.5 Mb that regulates the change in 6,699 CpGs (4.0% of all CpGs, P=8.5*10^-37^) and 455 genes (5.6% of all genes, P=3.1*10^-10^) in response to ISO. We observe that genes regulated by this hotspot (Figure 3A) are enriched for GO terms associated with cardiomyopathy (P=5.1*10^-11^) and mitochondria (P=5.1*10^-37^), among other significant terms (Figure 3B). Our initial candidate gene of interest for this hotspot was *Dicer1*, a crucial component of the RNAi machinery and which has a non-synonymous protein-modifying mutation in both the Balb and DBA2 backgrounds. Upon further inspection, however, we observed that this mutation had no effect on *Dicer1* gene expression (R=-0.11, P=0.53) nor did it correlate with changes in cardiac phenotypes such as heart weight (R=0.07, P=0.78), prompting us to look at other genes in the hotspot. The other major component of this hotspot is the *Serpina3* family of genes, generated due to a 14-fold gene duplication event in rodents that separated out the *SERPINA3* gene found in other mammals into *Serpina3a* through *Serpina3n*^48–50^. In humans, *SERPINA3* encodes for alpha 1-antichymotrypsin, a broad inhibitor of serine proteases with diverse roles across multiple organ systems and disease states^48^ and a proposed biomarker for doxorubicin-induced cardiac damage^51^. In rodents, however, these roles have been split between the many *Serpina3* isoforms. Our analysis of this hotspot specifically narrowed in on *Serpina3n*, the gene on the far 3’ end of the duplication event. *Serpina3n* contains 10 missense mutations on the Balb and DBA2 backgrounds that are not present on the B6, A or C3H backgrounds (Figure 3C). The first two of these mutations (G13R and C20S) are both predicted to be highly deleterious by PolyPhen2^52^ with scores of 0.994 and 0.888 out of a maximum possible deleterious score of 1.0. For the remaining 8 polymorphisms (T69K, L84V, V85M, K156R, T157A, S258F, L273M, R281K), we were able to explore their combined effects using DynaMut2^53^, which models the overall effects of multiple mutations on a deposited pdb structure (which, in the case of *Serpina3n,* did not include the first 20 residues and therefore could not take into account the two mutations predicted to be most deleterious). The remaining 8 AA changes, however, had a combined stability ΔΔG of 5.31, suggesting a significant loss of stability for the *Serpina3n* protein in the affected strains (Figure 3C). *Serpina3n* also possesses a strong *cis-*eQTL and very significant correlations with cardiac weight (R=0.57, P=3.9*10^-17^,Fig 3D), lung edema (R=0.42, P=7.4*10^-8^), fibrosis (R=0.44, P=5.5*10^-10^), and Left Ventricular Internal Dimension (LVID)(R=0.48, P=5.2*10^-12^) but not, notably, ejection fraction (R=-0.05, P=0.47). Additionally, we observed in a prior transcriptomics study^23^ that *Serpina3n* is present in a transcriptomic gene module that is strongly associated with changes in cardiac phenotypes in response to ISO. There is also a recent report from another group showing that knockout of *Serpina3n* significantly affects cardiac remodeling after a myocardial infarction^22^. Combined, these results gave us confidence that *Serpina3n* is the most likely candidate gene at this hotspot.

**Figure 3:**
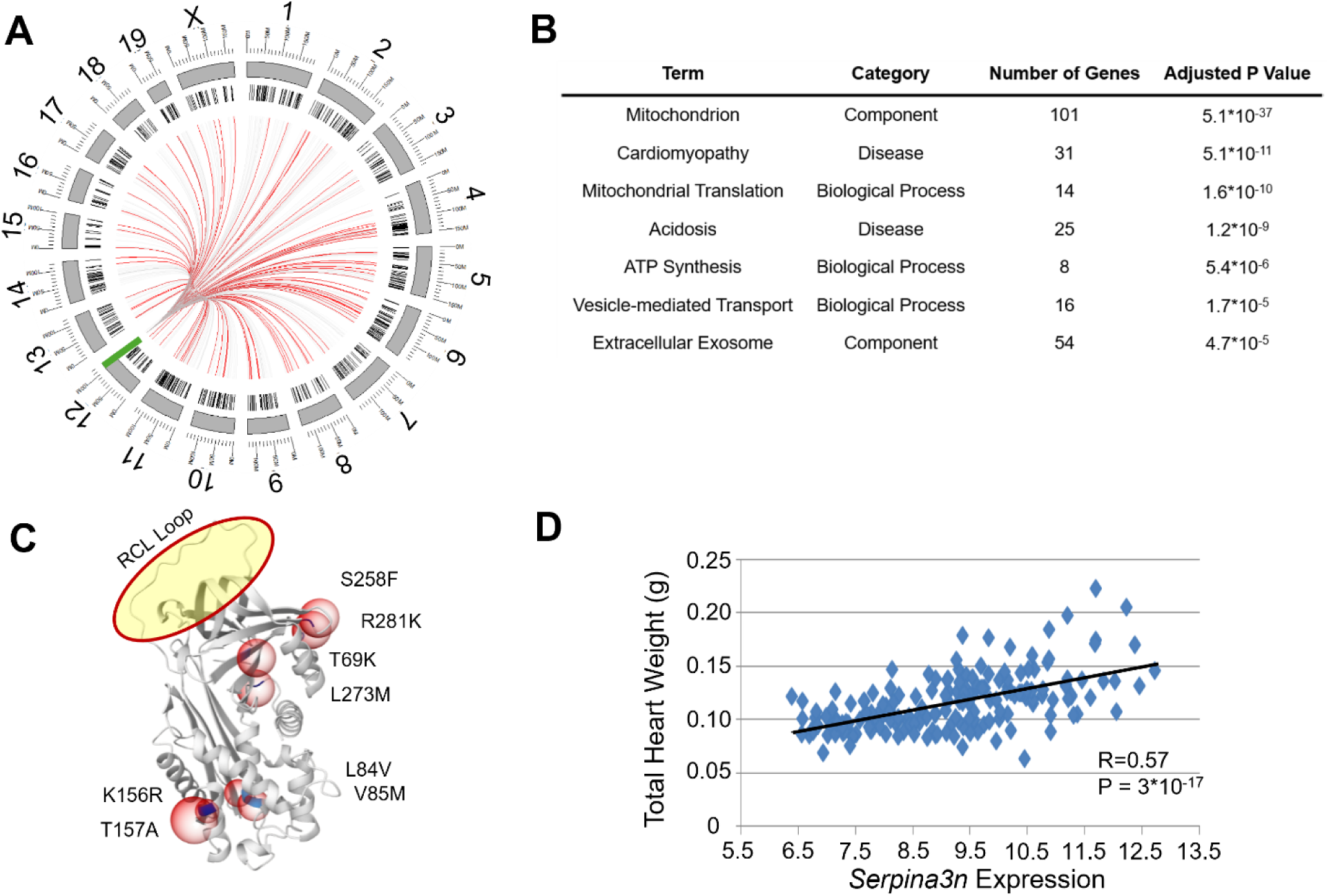
*Serpina3n* is the candidate gene for a hotspot on chromosome 12. A) Circular ideogram showing the associations between an eQTL hotspot on chromosome 12 (green bar) with 445 other genes (black bars). Grey lines indicate suggestive association (P<4.2*10^-5^) while red lines indicate significant associations (4.2*10^-6^). B) Selected enriched terms for genes regulated by this hotspot. C) Display of 8 of 10 structure-altering mutations in the *Serpina3n* molecule as rendered by MutationExplorer^87^. Red circles indicate mutation sites, while the yellow circle is the RCL Loop active site for the protein. Not displayed are G13R and C20S, the two mutations predicted to be most deleterious, that are located in a disordered region not present in the available structure. D) Correlation of *Serpina3n* expression with total heart weight across the HMDP panel

To test this hypothesis, we first examined whether siRNA-mediated knockdown of *Serpina3n* would result in changes to cardiomyocyte cross-sectional area (a cell-culture analogue of cardiac hypertrophy) in isolated NRVMs. We chose to test three possible hypertrophic stimuli - the beta-adrenergic agonist ISO, phenylephrine (PE), an alpha-adrenergic agonist, and angiotensin II (Ang II). We observe no significant effect of *Serpina3n* knockdown in untreated cells compared to a scramble control (P=0.24 and 0.88, respectively, for either *Serpina3n* siRNA used in this study)(Figure 4A). For each of the three hypertrophy-inducing treatments, however, *Serpina3n* knockdown significantly blunted their efficacy (Figure 4A). For example, while ISO treatment resulted in a 45% increase in average cell size compared with scramble control (P=2.4*10^-28^), cell sizes after siRNA-mediated knockdown only increased by 15-20% (P=1.1*10^-5^ and 1.2*10^-7^), reflecting an average 60.4% reduction in cardiac hypertrophy (P=2.5*10^-12^ and 7.1*10^-9^). Similar patterns and degrees of efficacy are seen in PE (average 56.3% reduction in hypertrophy, P= 1.7*10^-12^ and 3.9*10^-9^) and Ang II (average 53.4% reduction, P=1.8*10^-10^ and 8.6*10^-7^) (Figure 4A)

**Figure 4.**
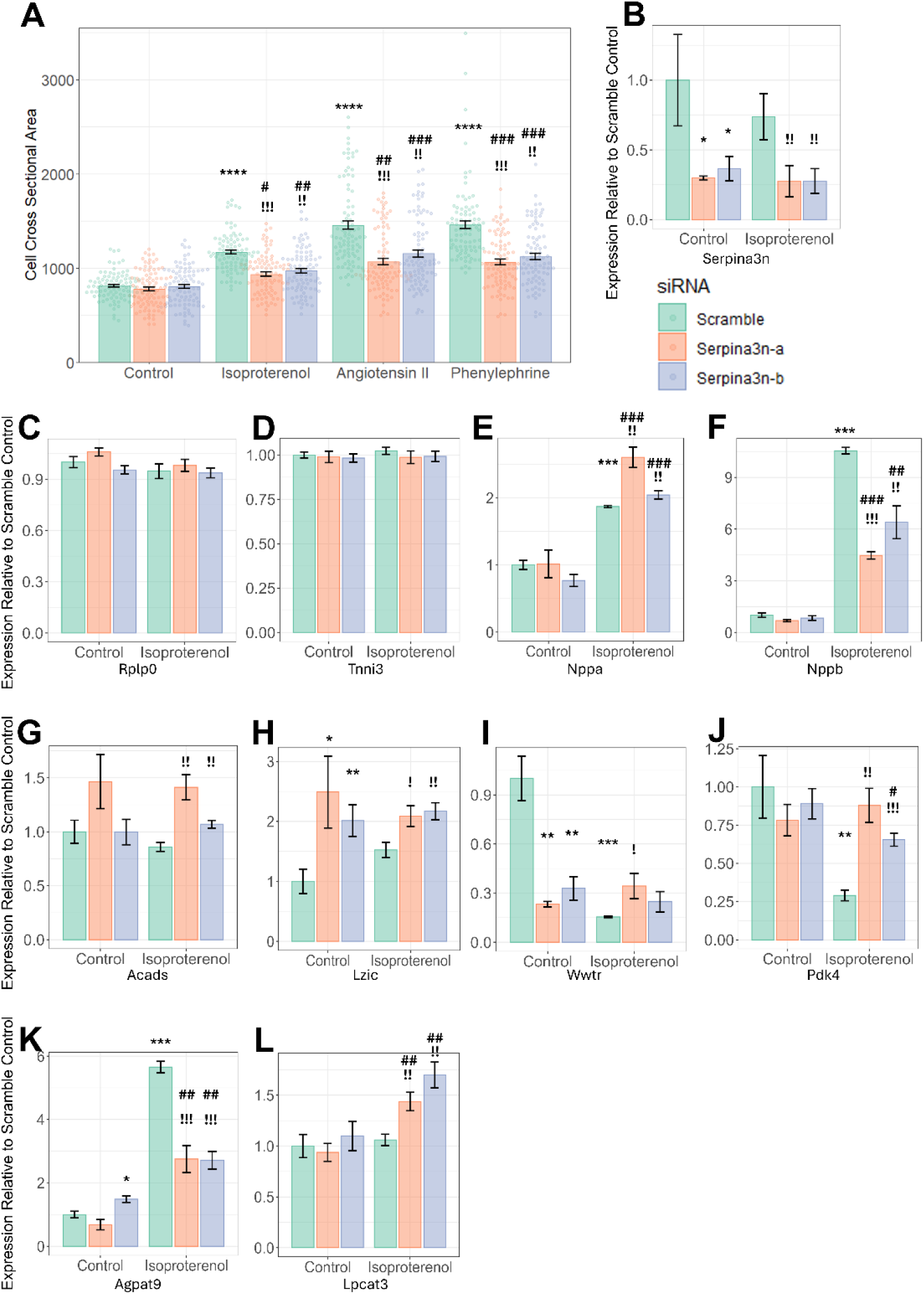
*in vitro* Analysis of *Serpina3n* knockdown in NRVMs. For all: Error bars indicate the SEM, * indicates significance vs control scramble (calculated for other control samples and scramble samples), # indicates significance vs respective control sample (e.g Isoproterenol Serpina3n-a vs. Control Serpina3n-a), ! indicates significance vs respective scramble sample (e.g. Isoproterenol Serpina3n-a vs Isoproterenol Scramble). Serpina3n-a and Serpina3n-b two different *siRNAs* used in this study A) NRVM cross-sectional areas for cells treated with Isoproterenol, Angiotensin II, or Phenylephrine and either a scramble siRNA or one of two *Serpina3n* siRNAs. * = P<.05, ** = P<1*10^-5^, ***=P<1*10^-10^, ****=P<1*10^-20^. N=90 per group, see Table S6 for full details B-L) Effects of *Serpina3n* knockdown on gene expression normalized to scramble control for B) Serpina3n, C+D) control genes Rplp0 and Tnni3, E+F) hypertrophy genes Nppa and Nppb and G-L) putative targets of *Serpina3n*: Acads, Lzic, Wwtr, Pdk4, Agpat9, Lpcat3. For all genes, *=P<0.05, **=P<0.005, ***=P<5*10^-5^. N=4-15 per group, see Table S7 for full details

We next examined the effect of *Serpina3n* knockdown on genes predicted to be highly affected by the hotspot through eQTL analyses. We selected the genes *Lzic (P=1.5*\*10^-10^*), Pdk4 (P=4.3*\*10^-9^*), Acads (P= 4.1*\*10^-8^*), Gpat3/Agpat9 (P=5*\*10^-8^*), Lpcat3 (P=1.3*\*10^-7^) and *Wwtr (P=1.6*\*10^-7^) as potential positive controls (P values are the eQTL significances for each gene at this hotspot), *Nppa* and *Nppb* as hypertrophic markers, and the housekeeping gene *Rplp0* and highly expressed but unaffected by ISO gene *Tnni3* as negative controls and performed qPCR analyses of their expression before and after ISO and with or without siRNA-mediated knockdown of *Serpina3n* in NRVMs. Both of our negative controls (Figure 4C,D), as expected, showed no change in expression due to either ISO or *Serpina3n* knockdown. Strikingly, in our hypertrophy markers, we observe differential dynamics between *Nppa* (Figure 4E) and *Nppb* (Figure 4F) expression. *Nppa* expression is unaffected by *Serpina3n* knockdown in untreated cells (P=0.88 and P=0.11, respectively), and shows an expected increase in scramble-siRNA cells after ISO stimulation (87% higher, P=1.4*10^-9^), yet after *Serpina3n* knockdown, despite reduced cardiomyocyte hypertrophy (Figure 4A) we observe maintained, or at times even significantly increased, levels of *Nppa* expression (2.5x higher, P=7.2*10^-8^ and 2.0x higher, P=1.7*10^-10^ compared to control scramble, 39% higher, P=5.2*10^-4^ and 9% higher, P=1.6*10^-2^ compared to ISO scramble). *Nppb* expression, on the other hand, matched our observed phenotypic effects, with an increased in *Nppb* expression due to ISO (12.0x higher, P=4.6*10^-12^), no change in untreated cells due to *Serpina3n* knockdown (P=0.80 and P=0.81), and a blunting of *Nppb* upregulation in *Serpina3n* knockdown cells (5.1x, P=1.9*10^-9^ and 6.6x, P=1.8*10^-4^), suggesting that *Serpina3n* may act downstream of the initial response to ISO, which stimulates the expression of *Nppa* and instead acts elsewhere, perhaps in the calcineurin-NFAT axis which modulates *Nppb* expression^22,54,55^.

For the genes expected to be affected by *Serpina3n* knockdown, in five of the six tested genes we observe that manipulation of *Serpina3n* levels leads to changes in cardiomyocyte size. The only gene which does not seem to significantly respond to *Serpina3n* knockdown is *Acads*, which shows a modest increase in average expression both before and after ISO in one siRNA (P=0.048 and P=0.017) but no effect in the other siRNA (Figure 4G). For two other genes, *Lzic* and *Wwtr,* we see that *Serpina3n* knockdown has mimicked the effects of ISO, with the addition of ISO having no further effect (Figure 4H, I). In one case, *Lzic* expression is increased with ISO in scramble cells (77% increase, P=8.4*10^-3^), but is upregulated at baseline in both siRNAs (2.9x, P=2.4*10^-4^ and 2.3x, P=2.6*10^-3^), with no further effect by ISO stimulation (P=0.24, P=0.61). Conversely, *Wwtr* expression drops after ISO stimulation in scramble siRNA cells (83% drop, P=6.1*10^-5^), with comparable losses in *Serpina3n* knockdowns at baseline (77%, P=1.4*10^-4^ and 67%, P=4.2*10^-4^), but no further loss of expression is observed with ISO (P=0.18, P=0.40).

For two other genes, *Serpina3n* knockdown blunts the modulation of the gene’s expression by ISO specifically. *Pdk4* (Figure 4J) drops by 70% in scramble ISO cells (P=4.6*10^-8^), but is unaffected by *Serpina3n* knockdown in untreated cells (P=0.37, P=0.65). When these knockdown cells are treated with ISO, one siRNA shows no effect of ISO on *Pdk4* expression (P=0.53), while the other shows a downregulation that is significantly weaker than that seen in the scramble-treated cells (24% decrease, P=0.045). Likewise, *Agpat9* expression (Figure 4K) increases 5.6x in scramble cells in response to ISO (P=1.3*10^-8^). One siRNA shows a comparatively modest increase in *Agpat9* expression at baseline (48% increase, P=3.5*10^-3^), while the other shows no significant change (P=0.16). In both siRNAs however, we see a blunting of the effect of ISO compared to WT (3.5x, P=6.7*10^-3^ and 1.8x, P=1.0*10^-3^). Finally, *Lpcat3* (Figure 4L) does not normally respond to ISO (P=0.63 in scramble ISO), and shows no effect by siRNA knockdown at baseline (P=0.67 and P=0.60), yet *Serpina3n* KO causes significant increases in *Lpcat3* expression after ISO (44% increase, P=3.2*10^-3^ and 70% increase, P=9.4*10^-3^).

## Discussion

This study combines data from prior research^4,23,30^ on isoproterenol-induced heart failure in the Hybrid Mouse Diversity Panel (HMDP), a genetic reference population comprised of both classical inbred and recombinant inbred mouse strains. Those studies used GWAS^4,10^ and EWAS^30^ approaches to link genetic and epigenetic mutations to changes in relevant cardiac phenotypes in response to beta adrenergic overdrive, a common pathway of HF exacerbation independent of initial etiology^56^. In contrast, here we combine these biological layers together to examine the effects of genetic variation on gene expression (expression QTLs or eQTLs) or CpG methylation sites (methylation QTLs or mQTLs), or how differences in CpG methylation can be associated with changes in transcripts (expression methylation-driven QTLs or emQTLs).

Of these three ‘flavors’ of cross-omics data, eQTLs have the longest history, first described by that name in the early 2000s^57,58^. Historically, a major use of eQTLs has been in understanding and refining the results of GWAS on phenotypic traits: searching for genes whose expression patterns were strongly correlated with the relevant phenotype to narrow down the list of possible candidate genes at a locus^59^. We have previously used eQTLs in this way in our GWAS and EWAS manuscripts^4,30^. Another application of eQTLs is to find loci that are responsible for the up/downregulation of a gene of interest in the context of a particular stressor^60^ or to better understand how changes in tissue or, more recently, cell type, interact with underlying genetic variation^61,62^. In a similar way, mQTLs have been used in the past to narrow down GWAS/EWAS loci^30,63^. These approaches are typically considered to be complementary or ancillary to the ‘main’ research focus of the GWAS or EWAS projects. The other major approach that uses eQTLs/mQTLs/emQTLs is to, instead of focusing on a single gene’s relationship with the genome, cast a wide net and look for genomic/epigenomic regions which regulate significantly more genes/CpGs than would be predicted by chance. These ‘hotspots’ are proposed to contain major regulators of the global gene/CpG landscape whose modulation could have outsized effects on phenotype through the combined effects of many affected genes^16,63–65^. We focused on these hotspots in this manuscript.

Each type of hotspot reflects a different sort of regulation. eQTL hotspots often reflect changes in transcription factors or chromatin modifiers, or the cascading effects of modulating the expression and/or processing of signaling molecules, leading to broad effects across the genome. Likewise, mQTL hotspots are caused by genes and mutations which drive changes in, among other things, DNA methyltransferases or methylases or, as with eQTLs, genes which regulate chromatin, as global shifts in methylation tend to follow shifts in histone marks^8^. Additionally, shifts in DNA methylation can modulate the expression of transcription factors or signaling molecules, driving emQTL hotspots.

We observe 204 unique (286 non-unique, 52 hotspots shared by at least two conditions) hotspots across our three analysis ‘flavors’ (eQTL: 49, mQTL: 98, emQTL: 139), although the larger size of the eQTL hotspots on average (1.3Mb) compared to mQTL hotspots (0.7Mb) means that the total amount of genome covered by eQTL and mQTL hotspots are comparable and a little over half that of the emQTLs (average hotspot size 0.9Mb). Although a smaller hotspot size in emQTLs is not surprising, likely tied both to the smaller ‘linkage disequilibrium’ of CpGs in the HMDP compared to SNPs^16,30^ and increased noise introduced into the EWAS formula by allowing continuous variation across both predictor and response variables vs GWAS where the SNP is discretized into 0, 1, or 2 minor alleles, it is noteworthy that eQTL hotspots are nearly twice the size of mQTL hotspots, given that both are regulated by the same sets of SNPs. This is possibly driven by the 10x higher number of variable CpGs analyzed vs expressed and varying genes in the genome.

When looking to validate our hotspots, we first set out to identify any hotspots that lined up with prior results from our GWAS or EWAS studies (see Table S4 for all such hotspots). We were able to identify 32 hotspots (15.7% of all hotspots) that overlap with hits from our prior studies (overlap P=1.0*10^-9^). The candidate genes within these hotspots reflect a variety of functions, from transcription factors (e.g. *Nfatc2*, long established as a key regulator of HF progression^66^), to transporters (e.g. *Abcc6*^4^ or *Slco3a1*^67^), to even some long non-coding RNAs implicated in cardiac pathologies (e.g. *Miat*^68,69^). Another six hotspots contain genes that were identified as hub genes in co-expression networks we generated from our transcriptome data^23^, including the metalloproteinase *Adamts2*, which we validated as a major regulator of a HF-driving co-expression module (in other words, like as in a hotspot, a gene whose manipulation results in the modulation of many other genes’ expressions), and *Serpina3n*, which we discuss in greater detail below. These overlaps not only provide evidence for hotspot validity, but also provide additional insight into the types of candidate genes to focus on when validating these GWAS/EWAS peaks.

We next focused our attention on hotspots whose effect on other genes of the genome are large– for example a delta eQTL hotspot on chromosome 1 that regulates 25.1% of the change in expressed gene abundances in the heart after ISO, and which contains *Phlpp1*, a gene linked to cardiac dysfunction^19^ whose action can be blocked by the flavonoid Isoquercitrin^18^. The outsized effects of these hotspots suggest that greater attention should be paid to the genes within them and their use as potential drug targets (e.g. *Phlpp1)*, as well as a case for follow-up studies to narrow down the candidate gene list in hotspots with multiple strong candidates, as is the case for an ISO eQTL hotspot on chromosome 8 that contains four reasonable candidates (*Ankr11, Spg7, Rpl13* and *Fanca)*.

Another option when looking for high-confidence hotspots is to, instead of looking for large numbers of transcripts modified by a locus, look for hotspots that appear across multiple analyses. Most of these hotspots appear within the same class of hotspot (e.g. one is ISO mQTL and the other is delta mQTL), cases where one result is an echo or reflection of another (a strong ISO peak, for instance, creating a weaker delta peak that is still detected). There are only nine hotspots in our study that are reflected across classes within their same condition (control eQTL and control mQTL, for instance), with only one hotspot present in all three classes which is located on chromosome 9. No gene is the clear candidate at this hotspot, although two, *Dync1li1* and *Cmtm8*, show strong correlations with cardiac phenotypes, most notably *Cmtm8*, which shows a strong correlation of its baseline gene expression not with control phenotypes, but with ISO phenotypes of hypertrophy (P=0.0155), edema (P=1.9*10^-4^), and ejection fraction (P=8.5*10^-5^). This mirrors what we have seen in our EWAS work^30^, where baseline DNA methylation was often highly predictive of treated phenotypes, likely representing a change to disease susceptibility through genetically-driven changes in gene expression that predate the addition of stressors. This has also been seen in other studies, for example where baseline expression of *KLRD1* in human NK cells was predictive of susceptibility to influenza^70^ and where differences in baseline immune cell types in the Collaborative Cross, another murine genetic reference population, was predictive of response to SARS-CoV infection^71^.

The most likely candidate gene in another one of these shared hotspots is *Serpina3n*, which has strong correlations to phenotypes such as cardiac hypertrophy (R=0.57, P=1.7*10^-15^), a highly significant *cis-eQTL*, and a series of highly deleterious amino acid changes in approximately 25% of the strains of the panel. We demonstrate that knockdown of this gene in NRVMs (Figure 4A) is capable of significantly blunting the hypertrophic effects of ISO and significantly modifying the expression patterns of five of the six tested genes predicted to be regulated by the hotspot via qPCR (the sixth being only suggestively (P<.05 but P>.01) predicted to be regulated by *Serpina3n*).

The Serpina3 family of genes is a 14-fold expansion of a single ancestral α1-antichymotrypsin that became SERPINA3 in humans, but in rodents became the Serpina3 family of *Serpina3a* through *Serpina3n*^22,48^. Although a number of the members of this family remain poorly described (e.g. 3a, 3f, 3h-m), others have recently been characterized as having organ or niche-specific roles. For instance, *Serpina3c* has been shown to ameliorate adipose tissue inflammation^72^ while *Serpina3k* deficiency reduces oxidative stress following kidney injury^73^ and *Serpina3k* lactylation protects against cardiac ischemia-reperfusion injury.^73^

*Serpina3n*, on the other hand, appears to have retained a broader scope similar to that seen in SERPINA3 in humans, with roles in the brain, where it appears to assist in maintaining learning and memory circuits^74^, the skin, where it improves the rate of wound healing in diabetic mice^75^, and systemically by blunting deleterious inflammatory responses in stroke, osteoarthritis, and other contexts^49,76^. In the heart, we first described a potential role for *Serpina3n* in a co-expression network derived from the HMDP heart transcriptome^23^, in which we observed that *Serpina3n* was a key member of a cluster of genes strongly correlated to cardiac hypertrophy and edema phenotypes^23^. Other groups have identified it as a biomarker for doxorubicin-induced cardiomyopathy^51^ and note that a similar role has been described for SERPINA3 in humans^50^. Recently, Sun et al^22^ compellingly demonstrated that *Serpina3n* plays an important role in regulating post-infarction remodeling, with significant upregulation in infarcted tissue. In their study, gene knockout of *Serpina3n* led to increased infarct size, cardiac fibrosis and inflammation, and proteolysis. In contrast, our study, in which we used a chronic stressor of either ISO, AngII, or PE, showed that knockdown of *Serpina3n* resulted in reduced cardiac hypertrophy while not affecting baseline cell sizes. This suggests that a key distinction between some of these other studies, such as the post MI^22^ or post traumatic brain injury or stroke^49,74^ models, and our own is the magnitude of activation of the inflammatory response – in the case of an infarction or a TBI, the damage is significant, severe, and immediate, whereas with catecholamine overdrive or AngII administration, the effects are chronic and cumulative over time. This suggests that modulation of the inflammatory pathways by *Serpina3n* may lead to context-specific benefits and disadvantages during acute or chronic cardiac stress. A possible place for further examination is the differences in regulation of *Nppa* vs *Nppb* expression after ISO stimulation with and without *Serpina3n* knockdown. As expected, in the scramble siRNA, we saw both *Nppa* and *Nppb* expression increase, yet after *Serpina3n* knockdown, we saw *Nppa* expression rise to a similar extent as seen in the scramble, but *Nppb* induction was significantly blunted. This points to *Serpina3n* playing a role in cardiomyocytes that is downstream of the initial adrenergic response to which *Nppa* is more sensitive but upstream of the Calcium/calcineurin to NFAT to GATA4 pathway that drives *Nppb* expression^77,78^. More research will be necessary to understand the exact mechanism of *Serpina3n*’s action in modulating the response to different cardiac stressors.

Our study has some limitations. First, our data comes from only female mice, which limits our ability to extend our results to male mice. During our pilot study, we noted larger variability in phenotype in response to ISO in the mice we tested and, due to cost constraints, chose to focus solely on female mice to maximize our statistical power. Although a limitation, past HMDP studies in both sexes have found that the vast majority of loci identified in one sex are observable in the other^11,32^. Another limitation is the differences in technology underlying our eQTL vs mQTL studies – our transcriptome data was drawn from RNA microarrays, which provide a continuous signal but struggle with issues of noise and oversaturation at the edges of their detectable dynamic range and which can limit analyses of high or low-expressed genes, especially at the same time. In contrast, our methylation data is drawn from reduced representational bisulfite sequencing, which is a discrete, counts-based approach with, theoretically, no limits on the dynamic range of the result. Comparing counts (RRBS or RNAseq) and continuous (microarray) data has received significant attention by researchers^79–81^ with no clear solutions. Consequently, beyond the different ‘omics levels involved, we were unable to use the same algorithms in each case, selecting a continuous-approach algorithm for our eQTL and mQTL data in EMMA and a counts-based algorithm for our emQTL data in MACAU. Another limitation of both RNA microarrays and RRBS is that they are representational approaches – neither captures the full breadth of expressed genes or methylated CpGs in the genome, although they are enriched in both cases for the most ‘relevant’ signals. This means that we may have missed some hotspots and seen some overall reduction in power that could be counteracted through RNAseq or whole genome bisufite sequencing approaches. Finally, our mQTL and emQTL hotspots tended to regulate a lower percentage of CpGs or gene transcripts than our eQTL results. This may be because of shifts in underlying cardiac cell type composition between samples that reduced power. As we described in our EWAS paper^30^, there are known shifts in cell type proportions as a response to ISO and underlying genetic variation that would affect detected CpG methylation values since CpG methylation is significantly different between cell types. These shifts in cell type are known confounders of EWAS studies (and, by extension, mQTL and emQTL studies)^82^ which are often addressed through the use of covariates when the cell type proportions are known (for instance, when the data is drawn from blood samples), but which are not feasible to do at a population scale in the heart. This uncertainty reduces the power of our mQTL and emQTL analyses and, consequently, the magnitude of the mQTL and emQTL hotspots. Approaches to deconvolute the bulk transcriptome data and estimate cell type proportions^83–86^ could be an initial step to counteracting this uncertainty and increasing the power of our analyses.

Cardiac homeostasis requires careful coordination of dozens of distinct biological pathways. Likewise, heart failure, a disruption of this homeostasis, involves the dysregulation of these pathways and the establishment of new, dysfunctional gene networks. Our results identify new avenues for analysis towards the identification of major regulators and coordination centers for these adaptive and maladaptive processes. By drawing data from across multiple ‘omics levels, we begin to shed light on trans-omics pathways that drive cardiovascular function and dysregulation. Further refinement of our hotspots and analyses of the genes contained within will allow for additional insights into the role of SNPs and CpGs in the progression of heart failure, with the ultimate goal of improved personal therapies for patients.

## Data Availability

RRBS data from the HMDP is available at the Sequence Read Archive at accession PRJNA947937. Gene Expression from the HMDP is available at the Gene Expression Omnibus at accession GSE48760. HMDP Phenotypic data is available through Mendeley Data at accession 10.17632/y8tdm4s7nh.1.

## Acknowledgements

This research was supported by NHLBI R00HL138301 and R01HL162636.

## Author Contributions

CL, SR, AR, CDR designed and performed the experiments. CL, AD, CDR performed the computational analysis. CL and CDR wrote the manuscript CDR supervised and conceptualized the project. All authors contributed to revising and editing the manuscript.

## Declaration of Generative AI and AI-assisted Technologies in the Writing Process

The authors did not use generative AI or AI-assisted technologies in the development of this manuscript.

