## Supplemental Information for "Global Analyses of Genomic and Epigenomic Influences on Gene Expression Reveals *Serpina3n* as a Major Regulator of Cardiac Gene Expression in Response to Catecholamine Challenge During Heart Failure"

**Supplemental Data**

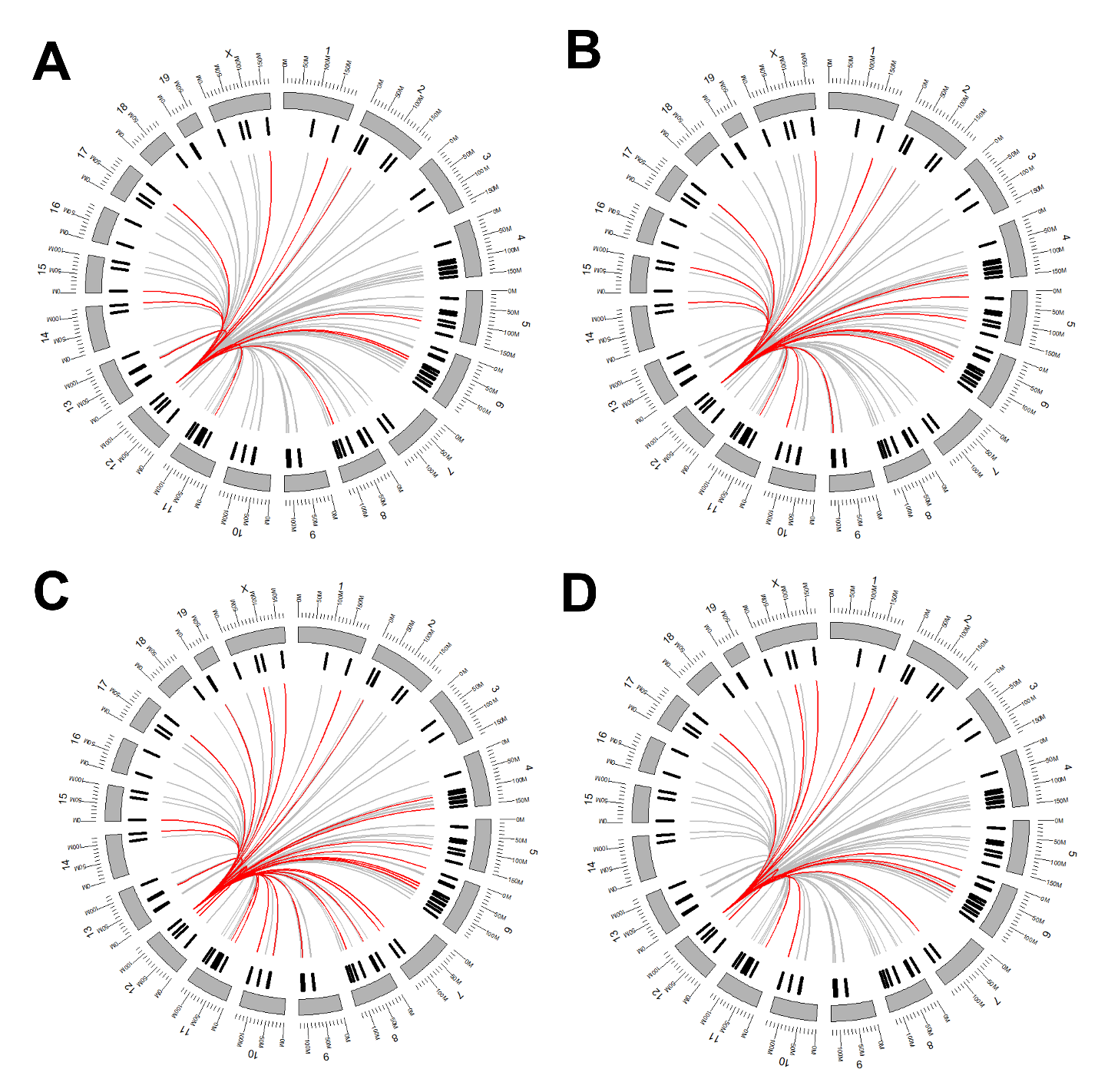

**Figure S1: Circle Ideograms for Selected GO Terms.** Ideograms generated for specific GO terms from the genome-wide significant genes linked to the *Serpina3n* locus on Chr12. A) Fatty Acid Oxidation B) Cardiomyopathy C) Mitochondrial Regulation D) ATP Synthesis

**Table S1:** **List of strains used in the study**

**Table S2: All sequences used in the study (primers + siRNAs)**

**Table S3: All significant peaks identified in the study**

**Table S4: Overlapping peaks with GWAS/EWAS or hypertrophic gene module results**

**Table S5: Enrichments of Genes associated with the *Serpina3n* hotspot**

**Table S6: All Values and Significances for Fig 4A**

**Table S7: All Values and Significances for Fig 4B-K**
